# Multimodal imaging for identifying brain markers of human prosocial behavior

**DOI:** 10.1101/2023.05.26.541897

**Authors:** Toru Ishihara, Hiroki Tanaka, Toko Kiyonari, Tetsuya Matsuda, Haruto Takagishi

## Abstract

How is the high degree of prosocial behavior that characterizes humans achieved? Here, we examined the structural and functional basis of the human brain with prosocial behavior using multimodal brain imaging data and 15 economic games. We identified that stronger interhemispheric connectivity, greater corpus callosum volume, higher functional segregation and integration, and fewer myelin maps combined with a thicker cortex were strongly associated with prosocial behavior. These associations were found especially in the social brain regions. This suggests that the strength of functional/structural connectivity between the left and right hemispheres, the strength of modular and efficient networks, and the high number of non-myelinated cells (i.e., dendrites, spines, synapses, and glia) are strongly associated with higher prosocial behavior in humans, particularly in the social brain regions.

## Main Text

Natural disasters, such as earthquakes and tsunamis, cause extensive loss of life and damage to the lives of residents. Numerous people from around the world have visited the impacted areas to help survivors, clear debris, and raise funds for reconstruction. Such prosocial behavior toward genetically unrelated individuals is one of the tendencies that characterize the human species relative to other animals (*1,2,3*). Human prosocial behaviors include altruism, such as volunteerism and charitable donations, fairness in resource distribution, cooperative behavior in collaboration, reciprocity of favors, and many other behaviors. The following questions then arise: why are humans more prosocial than other species and what are the critical factors that separate humans from other species? A possibility is that the function and structure of the human brain has evolved differently from that of other species. Although the human brain is not larger than that of other animals, the percentage of the neocortex in the brain is known to be extremely high (*4*). The proportion of the neocortex is related to the degree of social networking (*5*). In a social environment such as human society, which includes strangers, complex cognitive processing is undoubtedly necessary for building cooperative relationships with others. For example, the temporoparietal junction (TPJ) is the region involved in the cognitive ability to understand the mental state of others. This is called mentalizing, and higher-order mentalizing ability is prominent among humans (*6,7*). In addition to the TPJ, the temporal lobe, insular cortex, and inferior frontal gyrus are social brain regions (*8*). Many studies have revealed neural mechanisms behind human prosocial behavior, and it has been shown that neural activation in social brain regions forms the neural basis of human prosocial behavior (*9,10*). Despite the significant progress that has been made in understanding the neural mechanisms underlying human prosocial behavior, the full extent of the neural basis of prosocial behavior has yet to be fully elucidated. Since most previous studies have only examined relationships between single brain regions or relatively few regions utilizing limited neuroimaging methodologies, such as volumetric and activation techniques, further investigations are needed to fully understand the complex neural networks and processes that support human prosocial behavior. Considering the brain as a single system and elucidating its functioning for prosocial behavior are important for understanding the evolution of human prosocial behavior. In this study, we aimed to use data-driven analysis of multimodal magnetic resonance imaging (MRI) data to identify brain functions and structures underlying prosocial behavior.

Overall, 217 participants aged 20–60 years participated in 15 major economic games (explained in Supplementary Materials) and underwent MRI during 2012–2018. Participants played economic games with others in a completely anonymous environment and received rewards based on their actual behavior. Structural (T1- and T2-weighted), resting-state functional, and diffusion MRI data were collected and preprocessed using the Human Connectome Project pipeline tools (*11*) to obtain data for all 360 cortical and 41 subcortical brain regions, including data on cortical thickness, T1-weighted divided by T2-weighted (T1w/T2w) myelin maps, neurite orientation dispersion, neurite density, functional and structural connectivity based on graph theory indices (centrality, functional segregation, integration, and interhemispheric connection), and subcortical volume. A comprehensive analysis of the association between 108 behavioral data from 15 economic games and 5441 MRI data was conducted using the sparse multiple canonical correlation analysis (SMCCA) after dimension reduction using principal component analysis (PCA).

Economic games’ variates correlated with variates for cortical and subcortical structure (ρ = .47), functional (ρ = .35), and structural (ρ = .40) connectivity (nonparametric permutation test: *p* = 0.004; **Fig. S1**). Canonical cross-loading for economic games showed that prosocial behavior and aggression/punishment had positive and negative loadings, respectively, with brain imaging variates (**Fig. 1A**; **Table S1**). Canonical cross-loadings to economic games from brain imaging data consistently showed positive loadings of interhemispheric connections (interhemispheric functional connectivity: mean 0.08, 95% confidence interval [CI] 0.07 to 0.09, 90% regions showed positive loadings; interhemispheric structural connectivity: mean 0.04, 95% CI 0.03 to 0.05, 78% regions showed positive loadings; corpus callosum volume: mean 0.08, 95% CI 0.06 to 0.10, all regions showed positive loadings) (**Fig. 1B**; **Table S1**). Cortical T1w/T2w myelin maps had strong negative cross-loadings (Mean −0.08, 95% CI − 0.08 to − 0.07, 96% regions showed negative loadings) (**Fig. 1B**; **Table S1**). The positive cross-loadings were detected for cortical thickness (Mean 0.05, 95% CI 0.04 to 0.06, 72% regions showed positive loadings) and certain subcortical regions’ volume (nucleus accumbens, amygdala, hippocampus, and pallidum), while the ventricle showed negative loadings (mean −0.30, 95% CI −0.36 to −0.24, all regions showed negative loadings). The segregation measures of functional connectivity showed positive loadings (clustering coefficient: mean 0.04, 95% CI 0.03 to 0.05; local efficiency: mean 0.03, 95% CI 0.02 to 0.04; 72% and 63% regions showed positive loadings, respectively), while an integration measure (nodal path length) showed negative cross-loadings (functional connectivity: mean −0.04, 95% CI −0.04 to −0.03; structural connectivity: mean −0.05, 95% CI − 0.06 to −0.04; 71% and 77% regions showed negative loadings, respectively) (**Fig. 1B**; **Table S1**). Other measures did not show a consistent association with economic game variety.

**Fig. 1.**
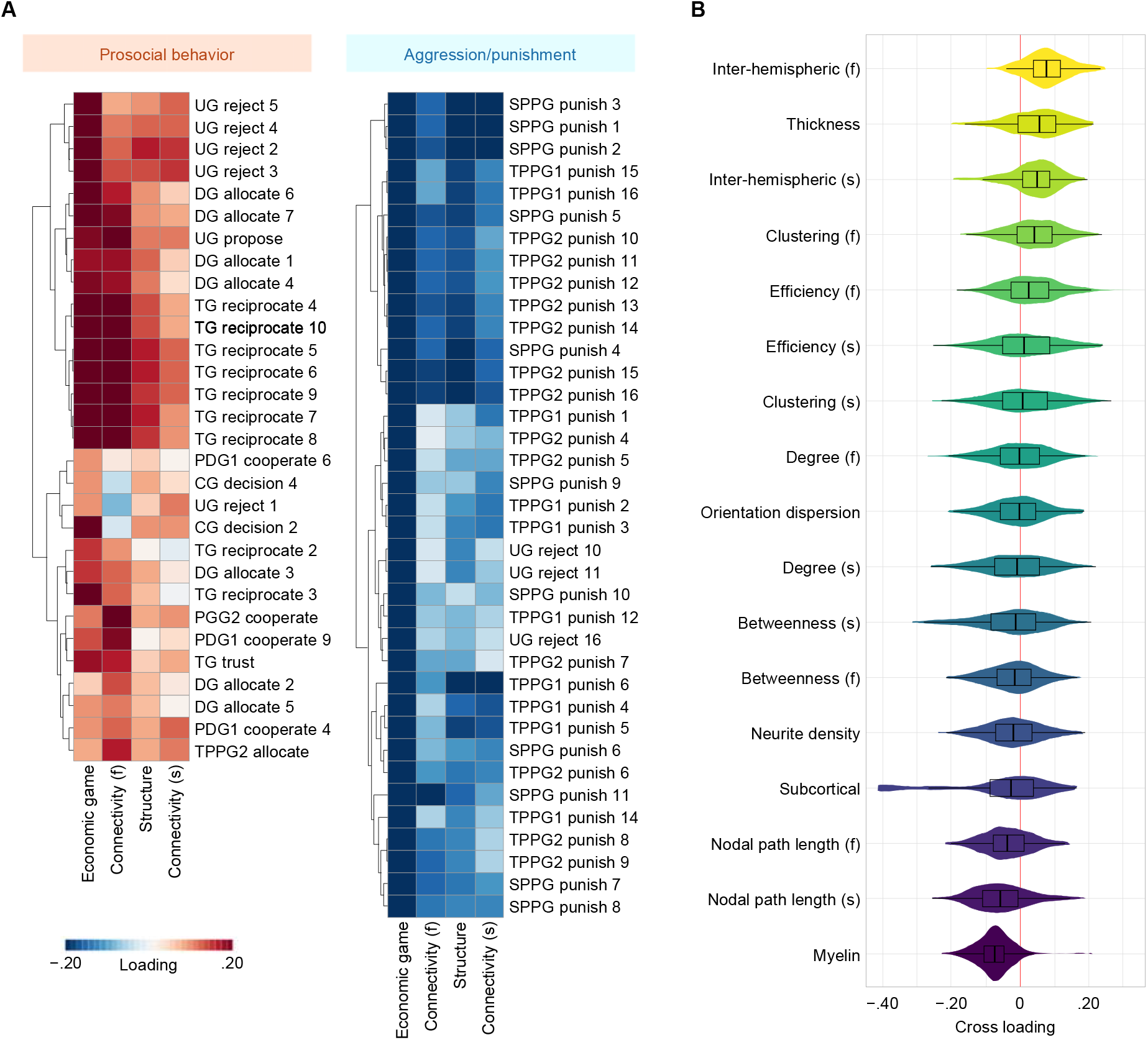
Results of multiple sparse canonical correlation analysis. (**A**) Canonical cross-loadings for prosocial behavior. Canonical loadings for prosocial behaviors are presented only in clusters with consistent positive loadings (left panel) and negative loadings (right panel), which reflect prosocial behavior and aggression/punishment, respectively. The full results are presented in **Fig. S2**. (**B**) Mean canonical cross-loadings for brain imaging data. Canonical cross-loadings of the MRI dataset (5441 images) are divided into each MRI metric and presented as violin and box plots. (f) denotes measures of resting-state functional connectivity; (s) denotes measures of tractography-based structural connectivity. Descriptions of the variables measured in each economic game are presented in **Table S2**. UG, ultimatum game; DG, dictator game; TG, trust game; PDG, prisoner’s dilemma game; CG, chicken game; PGG, public goods game; TPP, third-party punishment game; SPP, second-party punishment game.

The inter-region variations of canonical cross-loading are presented in **Fig. 2**. As the cortical thickness must be considered for the interpretation of individual differences in cortical T1w/T2w myelin maps (*12*), a correlation analysis was performed. The cross-loading of T1w/T2w myelin maps inversely correlated with that of cortical thickness (Pearson’s r −.23, 95% CI −.32 to −.13). When the imaging data were summarized according to brain regions that most strongly contribute to the prosocial behavior (top 30%), MRI data strongly covaried with economic games (i.e., cortical thickness, T1w/T2w myelin maps, nodal path length, and interhemispheric connectivity) a pattern emerged (**Fig. 3**) that included the temporal, parietal, insula, and inferior frontal regions, which are well known as social brain networks. However, no such specific patterns were seen in other lest of MRI indications (**Fig. S3**). The pre-SMCCA dimension reduction using PCA was run using a variable number of PCA, with almost no change in the results—original versus alternative correlations [0.76 to 0.99] (**Figs. S4** and **S5**).

**Fig. 2.**
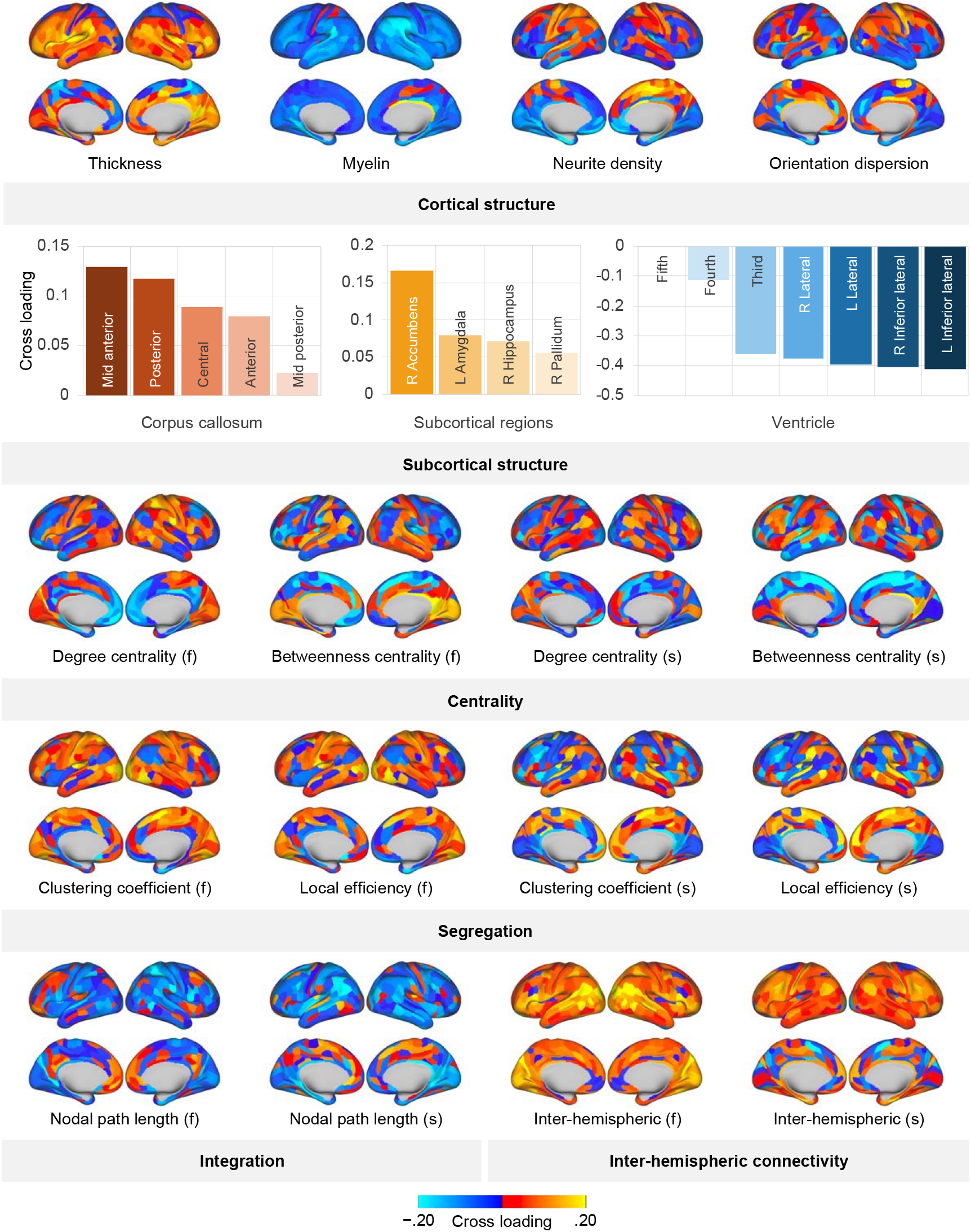
Canonical cross-loadings for brain imaging data in each region. Only the canonical cross-loading most strongly associated with the economic games variate is presented for subcortical volume (for more quantitative results, please refer to **Table S1**). (f) denotes measures of resting-state functional connectivity; (s) denotes measures of tractography-based structural connectivity.

**Fig. 3.**
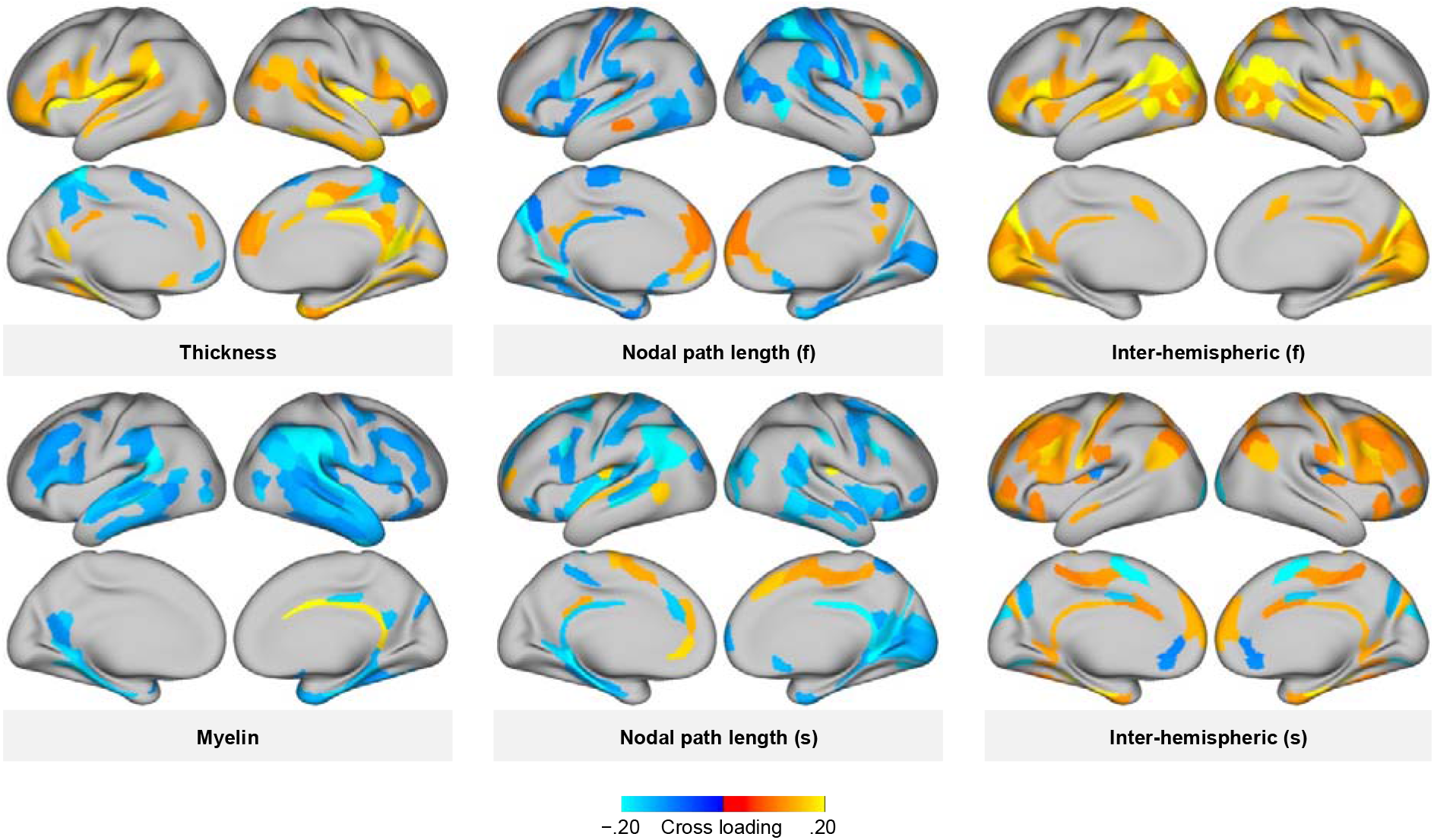
A brain regional pattern of the strong covariation of brain imaging data with economic games. This figure maps brain regions that show only the top 30% canonical correlation coefficients to confirm the brain regions strongly associated with prosociality. (f) denotes measures of resting-state functional connectivity; (s) denotes measures of tractography-based structural connectivity.

The comprehensive data-driven analysis using SMCCA identified brain functions and structures that support prosocial behavior. The higher the degree of prosocial behavior, the greater was the functional connectivity between cerebral hemispheres. The results indicate that during the resting-state brain function, the degree of prosocial behavior is higher in individuals with a better functional balance in the same brain region between the cerebral hemispheres. Additionally, individuals with stronger structural connectivity between hemispheres and those with greater corpus callosum volumes exhibited higher prosocial behavior. These findings are consistent with previous research indicating that patients with autism spectrum disorders, which are primarily characterized by social deficits, showing weaker resting-state functional- and tractography-based structural connections between hemispheres (*13,14*) and smaller volumes of corpus callosum (*15*). The present results of the association of decreased ventricular volume with a lower degree of prosocial behavior accord well with those of a previous study suggesting that patients with autism spectrum disorders have larger ventricular volumes (*16*). The corpus callosum is a collection of white matter fibers that connect the left and right hemispheres. Compared with chimpanzees, humans have a larger volume of the rostral body of the corpus callosum, which is involved in behavioral control (*17*), and humans exhibit higher prosocial behavior than chimpanzees (*18,19*). Altogether, this evidence suggests that the evolution of the interhemispheric connections supported by structural connectivity and the large volume of the corpus callosum might be a possible answer to why humans are more prosocial than other animals, and what are the critical factors that separate humans from other animals.

Following the interhemispheric functional and structural connectivity, decreased T1w/T2w myelin maps were also associated with a higher degree of prosocial behavior. Myelin is an insulator in the axon of a neuron and is involved in the efficiency of neurotransmission (*20,21*). Generally, lower T1w/T2w myelin maps are interpreted as demyelination in the axon; however, the association of T1w/T2w myelin maps with prosocial behavior should be interpreted with the results of cortical thickness. The present results demonstrated that a higher degree of prosocial behavior is associated with a thicker cortex. Decreased myelination simultaneous with increased cortical thickness may be due to increased non-myelinated plasticity supporting cellular constituents, such as dendrites, spines, synapses, and glia, and not necessarily due to an absolute decrease in myelination (*12*). Therefore, the present findings suggest that one or more of these non-myelinated constituents may play a significant role in promoting human prosocial behavior. The covariation of interhemispheric connectivity, T1w/T2w myelin maps, and cortical thickness with the prosocial behavior were found in the whole brain; however, it was more pronounced in the social brain network (*8*), including the temporal lobe, TPJ, insula, and inferior frontal gyrus. These findings support the previous line of literature showing the prominent role of social brain networks on human prosocial behavior (*9,10*). Previous studies on cortical thickness have found that people with thicker cortical thickness have lower allocation in the dictator game (*22*), suggesting that the dorsolateral prefrontal cortex (DLPFC) inhibits prosocial impulses. As this study uses some of the same data as those used in a previous study (*22*), at first glance, the results of the present study may appear to be contradictory to those of the aforementioned previous study. However, in this study, the prosocial component was extracted from various economic games, not only the dictator game, and the relationship between various MRI indices in the whole brain was examined. In addition, unlike this study, previous studies focused analysis on the DLPFC and did not report any association with cortical thickness in the social brain regions, including the TPJ, which was found to be relevant in the present study. In the brain, the association between prosocial behavior and DLPFC was relatively weak, and not necessarily contradictory, as it was more closely related to the involvement of social brain regions, such as the TPJ and insula, as described above.

Several graph theory indicators also showed associations with prosocial behavior. First, a higher clustering coefficient and local efficiency, as well as a shorter nodal path length, were associated with a higher degree of prosocial behavior. These results indicate that the higher modular segregation and the quicker inter-regional communications, the higher the prosocial behavior, in other words, the more efficient it is. The association between integration and prosocial behavior is prominent in the social brain regions, while no such pattern was found for segregation, suggesting that human prosociality is supported by a social brain network that allows rapid communication with other regions. Although results that suggest the relation of TPJ to prosocial behavior have been previously reported (*9,10*), this study contributes majorly to show that high prosocial behavior is achieved by rapid communication of social brain-related regions including the TPJ.

This study used multimodal brain measures and many major economic game measures to identify brain functions and structural bases underlying prosocial behavior in over 200 adults. This approach, called population neuroscience (*23*), uses a large amount of MRI data of many subjects, and we believe that it is effective for obtaining more robust results. However, dealing with large amounts of data involving multisets of multiple variables can present a disadvantage as the interpretation becomes complex if using repeated analyses, such as one-to-one or many-to-one relationships like Pearson’s correlation and multiple regression. Furthermore, repeated analysis should be avoided due to the risks of type 1 and type 2 errors. To address these analytical issues, the SMCCA approach presented in this study is effective as it allows the identification of patterns that describe many-to-many relationships. Based on the present results obtained by the use of these approaches, we conclude that the neural basis of human prosocial behavior involves strengthened interhemispheric connections, a larger cortex due to increased non-myelinated tissue, and functional segregation and integration in the social brain network. The study provides evidence that the human brain is uniquely adapted to support prosocial behavior, which may have contributed to the evolution of human sociality.

## Acknowledgments

We thank Atsushi Miyazaki, Takayuki Fujii, Muneyoshi Takahashi, Kuniyuki Nishina, Kei Kanari, Chiaki Ishiguro, Yoshie Matsumoto, Yang Li, Masamichi Sakagami, and Toshio Yamagishi for their help in conducting the study.

## Author contributions

Conceptualization: TI, HTan, TM, Htak

Data curation: TI, Htan, Htak

Formal analysis: TI

Methodology: TI

Investigation: TI, Htan, Htak

Visualization: TI

Supervision: TK, TM

Project administration: TM, Htak

Funding acquisition: Htak

Writing–original draft: TI, Htak

Writing–review & editing: Htan, TK, TM

## Data and materials availability

Behavioral data are available at https://osf.io/r2ewv/?view_only=f6bb9247afa74713a23337a24c280435

## Supplementary Materials

Materials and Methods

Figs. S1 to S5

Tables S1 to S4

