## Supplementary material for "Multimodal imaging for identifying brain markers of human prosocial behavior": Materials and Methods

Participants

This paper is a secondary analysis of data collected in an earlier research project (Neuropsychological and Social Institutional Foundations for Prosocial Behavior http://www.human-sociality.net/english/). Approximately 400 men and women in their 20s to 60s and living in Tokyo participated in repeated experiments from 2012 to 2018. They participated in 10 experiments during this period, where they played various economic games and performed cognitive tasks, answered psychological questionnaires, and underwent MRI imaging and collection of saliva and buccal cell samples. Experiments were conducted multiple times to reduce the burden on participants and overcarry effect in the economic game. The schedule for each experiment is shown in **Table S3**. To avoid arbitrary economic game selection, the analysis was conducted using behavioral data from all economic games collected in the project. However, as the number of participants decreased after the tenth wave, the analysis was conducted only for the economic games conducted in the second through eighth waves. The experimental protocol for this study was approved by the Tamagawa University Ethics Committee (approval number: TRE18-030), and all participants completed a consent form before participating in each experiment. Of all participants, 217 who participated in all economic games and underwent MRIs were included in the analysis. The data collected in this project have been reported in many other papers (**Table S4**). However, this paper is the first to conduct a comprehensive analysis using data from all economic games and MRI scans.

Economic games

All economic games were performed in a completely anonymous situation. Each session included several participants simultaneously; each participant worked in an individual booth. All experiments did not use deception and participants received rewards based on their actual behavior. Descriptions of the variables measured in each economic game and used in the analysis are presented in **Table S2**.

1. *Prisoner's Dilemma Game I (PDG I)*

The PDG-I was conducted in pairs. Participants chose between providing the endowment received from the experimenter to their opponent or not. The endowment offered to the opponent was doubled and given to the other player. Participants underwent nine PD trials with a different partner each time. There were three experimental conditions in this task: a situation in which two participants made decisions at the same time (simultaneous PDG); a situation in which the participants made a decision first and the opponent made a decision later (sequential PDG first), and a situation in which the opponent made a decision first and participants decided later (sequential PDG second). These three situations were presented to participants three times each in random order. In the case of sequential PDG second, they answered how they would decide if their partner offered or did not offer using the strategy method. The endowment received from the experimenter was either JPY 300, JPY 800, or JPY 1500, and was provided to participants in random order.

2. *Dictator Game (DG)*

The DG was conducted in pairs, with one participant in the role of a dictator and the other in the role of a recipient. Participants initially performed one-shot DG. All participants were first assigned to the role of the dictator and were asked to decide the amount they would allocate JPY 1000 to their partner. After the decision was made, the participant's role and partner were randomly determined, and the reward amount was determined based on the results of the participant's actual behavior. After the game, participants were told to perform repeated one-shot DG. Participants then played the dictator game six times, changing opponents each time they played as a dictator. The endowment given to the dictator could be JPY 300, JPY 400, JPY 600, JPY 700, JPY 1200, or JPY 1300, and was offered in random order.

3. *Faith Game (FG)*

The FG was played in pairs. In this game, as in DG, there are two roles: the distributor, who is responsible for distributing JPY 1000 received from the experimenter, and the recipient, who is responsible for receiving the distributed money. The money distributed by the distributor was tripled and given to the recipient. Participants decided the amount of the endowed JPY 1000 they were certain to receive and the amount that they were willing to leave to the distributor for distribution and receive as a recipient. If the participant thought that the distributor would distribute more money to them, the participant would give more money to the distributor. Conversely, if the participant did not think that the distributor would distribute more money to them, then they decided to ensure that they receive more money.

4. *Public Goods Game I (PGG I)*

The PGG-I was played with several participants divided into groups of at least four or more. They decided in increments of 100 whether they were willing to contribute the JPY 1000 which they received from the experimenter for the group. Money contributed to the group was tripled and distributed equally among all participants. The experiment was completed in one trial.

5. *Prisoner's Dilemma Game II (PDG II)*

The PDG-II is played in pairs. Participants decided the amount of the endowed JPY 1000 they would provide to their partner, in increments of JPY 100. The endowment offered was doubled and given to the opponent. The opponent was also determined at the same time. The experiment was completed in one trial.

6. *Second-party punishment game (SPPG)*

Participants played a one-shot, two-person SPPG, consisting of two phases—the PDG phase and the punishment phase. In the PDG phase, participants were randomly matched with another participant, and they played a PDG with the opponent, knowing that both players would have a chance to reduce the other player's earnings by spending some of their own money, up to an amount of JPY 1500. In the PDG phase of the SPPG (PD/SPPG), each player was endowed with JPY 1000 and was instructed to decide the amount of the endowed money they were willing to give to their opponents (in increments of JPY 100). The money given by the player was doubled and was transferred over to the opponent. The task was symmetrical, such that the player would receive twice the amount of money his/her opponent gave to him/her as well. Therefore, each player received the portion of the endowment that he/she did not give, plus twice the amount of money his/her opponent gave. When participants played this game, they knew that they would both have a chance to subtract money from the other player's cumulative earnings.

When the PD/SPPG was over, each participant was informed of their opponent's choice and were then asked to decide the amount (up to JPY 1500, in increments of 100) they would spend to reduce their opponent's earnings. The strategy method was used to determine the expenditure for punishment; that is, each participant was provided with a list of possible opponent cooperation levels, ranging from JPY 0 to JPY 1000, in increments of 100, and each was asked to decide the amount to spend in each case. No extra money was provided specifically to use for this spending. The money came from their earnings in earlier tasks and the show-up fee, although, at that time, the participants were not informed about of their accumulated earnings amount. The amount spent was doubled, and this new amount was then subtracted from the total earnings (including earnings from other tasks) of the opponent. Participants were explicitly told that the amount of money taken away from their opponents would not be transferred to them.

7. *Trust Game (TG)*

The TG was played by two persons, a trustor and a trustee. The trustor was asked to decide the amount of the endowed JPY 1000 to be transferred to the trustee in increments of 100. The amount of money transferred was tripled and received by the trustee. Next, the trustee was asked to divide the money between himself and the other participant. The decisions of the trustee were analyzed using the strategy method, which determined how the trustee would return to every decision the trustor could make, in increments of 10%. All participants in the experiment first made decisions as a trustor, then changed partners and made decisions as trustees. After the decision as a trustee, the game was over.

8. *Ultimatum Game (UG)*

The UG was conducted in pairs: the proposer and the responder. First, the proposer received JPY 1500 from the experimenter and proposed a plan to divide it between himself and the responder in increments of 100. After the proposer's decision was made, the responder decided whether to accept or reject the proposal. If the responder accepted the proposal, both participants received the money as proposed. In case of rejection of the proposal, neither received anything. In the experiment, all participants first took the role of the proposer, followed by the change of partners and decision as a responder. The responder's decision was made using the strategy method, i.e., acceptance or rejection of all proposals that the proposer could make (from JPY 0 to JPY 1500). The experiment was over when the decision as a responder was completed.

9. *Third-party Punishment Game I (TPPG I)*

The TPPG-I was played by three players. First, the distributor distributed the JPY 1500 received from the experimenter between himself and the recipient in 100 increments. The recipient received the amount decided by the distributor. After the distributor's decision, a third-party decided whether to reduce the distributor's money. Three times the amount of money spent by the third-party was deducted from the distributor's amount. The third-party could spend between JPY 0 and JPY 500 using money earned in another experiment. In the experiment, participants first took the role of the third-party and used the strategy method to decide the amount they wanted to spend to punish the distributor for all decisions that he or she could take. After the third-party decisions were made, the next trial began, and three new participants were matched. The participants took the role of distributor and decided how to distribute the JPY 1500 between themselves and the recipients, following which another three new people were matched; then the participant took the role of a recipient and received the money distributed by the distributor. The experiment ended there.

*10. Third-party Punishment Game II (TPPG II)*

TPPG-II differed from TPPG-I as the source used by the third-party to deduct the distributor's money was the amount received from the experimenter (JPY 500). All other procedures were identical.

*11. Preemptive strike game (PSG)*

The PSG was played in pairs (player A and player B). The buttons were displayed on each other's personal computers (PCs). If no one pressed the button for 30 seconds, both players received JPY 1500. However, if player B pressed the button first, player A received JPY 500, and player B received JPY 1300, and vice versa. If either one of the players pressed the button, it showed up on the screen indicating that the other player would not be able to press the button. In other words, only one of the players could press the button. The experiment involved four conditions in which both parties received different amounts of money, which were presented to the participants in random order. The matching partner changed each time. Condition 55: As described above. Condition 1010: if player B pressed the button first, player A received JPY 1000, and player B received JPY 1300. If player A pressed the button first, player A received JPY 1300, and player B received JPY 100. Condition 510: if player B pressed the button first, player A received JPY 500, and player B received JPY 1300. If player A pressed the button first, player A received JPY 1300, and player B received JPY 1000. Condition 105: if player B pressed the button first, player A received JPY 1000, and player B received JPY 1300. If player A pressed the button first, player A received JPY 1300, and player B received JPY 500.

*12. Chicken Game (CG)*

CG was a game played in pairs. Participants could choose to either "proceed" or "turn back" simultaneously. If both participants decided to "proceed", they received JPY 0. If both participants chose to “turn back,” the amount of money both participants received was JPY 300. If one chose to “proceed” and the other chose to “turn back,” the one who chose to “proceed” received JPY 1200, and the one who chose to “turn back” received JPY 300. The game was played thrice, each time with a different opponent, and three different amounts were received based on the decisions of both players. These three payoff matrices were presented to the participants in random order.

Payoff matrix 1.

| | | Player B | |
|---|---|---|---|
| | | proceed | turn back |
| Player A | proceed | 0 / 0 | 1200 / 300 |
| | turn back | 300 / 1200 | 300 / 300 |

Payoff matrix 2.

| | | Player B | |
|---|---|---|---|
| | | proceed | turn back |
| Player A | proceed | 0 / 0 | 1200 / 100 |

| | | | |
|---|---|---|---|
| | turn back | 100 / 1200 | 100 / 100 |

Payoff matrix 3.

| | | Player B | |
|---|---|---|---|
| | | proceed | turn back |
| Player A | proceed | 0 / 0 | 400 / 100 |
| | turn back | 100 / 400 | 100 / 100 |

Next, the participants were paired with a new partner. They played a game in which the participant decided first, and the opponent decided later (participant first condition). In this case, payoff matrix 1 was used as the reward combination.

*13. Public Goods Game II (PGG II) and Public Goods Game with Punishment (PGGW)*

Participants played a one-shot public goods game (PGG-II). The rules of the game were the same as those of PGG-I. Next, the participants were told that they would be playing a one-shot public goods game with punishment. In this game, the player who offered the lowest amount in the public goods game was punished. The amount for punishments were deducted from the funds for the punishment system, which was tallied by each player. Twice the sum of the amounts was deducted from the player with the lowest contribution. Each player could contribute between JPY 0 and JPY 300 in increments of 10 from the rewards earned in the experiment to fund the punishment system. The game ended when the amount of money for punishment was determined.

*14. Stag Hunt Game (SHG)*

The SHG was conducted in pairs. Participants decided whether to invest the JPY 500 they received from the experimenter or keep it with them. If both participants decided to invest, they received JPY 1000. If both decided to keep it in hand, both received JPY 500. If one participant decided to invest and the other decided to keep the money, the one who decided to invest the money received JPY 0, while the participant who decided to keep the money received JPY 500. Both decisions were made simultaneously, and the game ended with a single decision.

MRI data acquisition, preprocessing, and analysis

Using a 3-T Trio Tim scanner (Siemens, Erlangen, Germany) equipped with a 32-channel head coil, all MR images (T1- and T2-weighted, resting-state functional MRI (fMRI), and diffusion-weighted (dMRI)) were collected from November 11, 2016, to March 3, 2018, at Tamagawa University Brain Science Institute (Machida, Japan). The parameters used in resting-state fMRI and dMRI acquisition are as described elsewhere (*1*). The resting-state fMRI data and diffusion-weighted images were acquired twice with reversed phase-encoding direction (anterior-posterior and posterior-anterior).

In the preprocessing and analysis of MRI data, the Functional Magnetic Resonance Imaging of the Brain (FMRIB) Software Library (FSL; version 5.0.9), FMRIB's ICA-based Xnoiseifier (FIX; version 1.062), FreeSurfer (version 5.3.0-HCP), Human Connectome Project Pipeline (version 3.22.0), and ConnectomeWorkbench (version 1.2.3) were used. The Human Connectome Project structural preprocessing pipelines (*2*) which include PreFreeSurfer,

FreeSurfer, and PostFreeSurfer components were used to analyze cortical structure. The PreFreeSurfer pipeline was utilized to 1) align and average repeated T1- and T2-weighted scans of good or excellent quality (when available); 2) create an unbiased "native" volume space for each participant that is rigidly aligned to the Montreal Neurological Institute (MNI) template by removing gradient nonlinearity and readout distortion (static field [b0] distortion in three-dimensional images) to; 3) perform cross-modal alignment between the T1- and T2-weighted images by the use of FreeSurfer's boundary-based registration (BBR) method (*3*); 4) perform bias field correction with the square root (T1-weighted × T2-weighted); and 5) perform non-linear volume-based registration to the MNI template using FSL's FNIRT algorithm.

A customized version of FreeSurfer version 5.3 recon-all was used to generate white and pial cortical surfaces (using both T1- and T2-weighted volumes at 0.8-mm resolution), including subcortical segmentation, all carried out in the participants' native volume space. PostFreeSurfer was used to convert the FreeSurfer data into standard NIFTI, GIFTI, and CIFTI file formats and bring the data into MNI space. To perform an initial, gentle, non-rigid surface registration based on folding patterns (MSMSulc), the Multimodal Surface Matching (MSM) surface registration algorithm (*4*) was used. This technique has taken the place of the FreeSurfer folding-based registration utilized in earlier studies (*2*) because it yields slightly better initial alignment of functionally corresponding regions (such as task fMRI) while producing significantly less local distortion than the FreeSurfer algorithm (*4*). This registration together with the FNIRT non-linear registration was used to bring an initial version of the data into standard grayordinate space (32-k standard mesh for each hemisphere's cortical surface at 2-mm average vertex spacing and 2-mm isotropic MNI-space voxels for the subcortical volume data). The FreeSurfer-generated measure of cortical thickness was corrected for folding-related bias by regressing out the FreeSurfer mean curvature measure from each participant's thickness data (*5*) because the gyral crowns tend to be thicker than the sulcal fundi. The ratio of T1-/T2-weighted images and normalized for residual transmit field inhomogeneity were used to compute myelin maps (*2*, *4*–*6*).

All resting-state fMRI data were analyzed using the HCP functional preprocessing pipelines including volumetric- (fMRIVolume) and surface-based (fMRISurface) components (*2*, *7*). The fMRIVolume pipeline consisted of the following steps: removing gradient non-linearity and b0 inhomogeneity-related image distortions; motion correction; cross-modal alignment to the T1-weighted image with BBR (*3*); concatenation of all transforms, including the non-linear volume registration to MNI space; and resampling the original timeseries into MNI space using a single spline interpolation. A number of intensity normalization steps were used, including a crude fMRI bias field-correction step based on the structural data from a separate imaging session (this bias field correction is changed to a better method in the following processing steps, as described below, and has been incorporated into the most recent version of the pipelines), and grand four-dimensional mean normalization to 10,000. Then, to map gray matter timeseries data into the 91,282-grayordinate standard space (2-mm average cortical vertex spacing and 2-mm subcortical voxels) with a 2-mm full-width at half-maximum smoothing kernel (constrained to the cortical surface and subcortical gray matter segmentation), fMRISurface pipeline was utilized. For each resting-state fMRI run, these steps produce a "dense timeseries" CIFTI file. To replace the crude bias field correction map with a better map, it was also mapped into standard CIFTI space (by dividing it back and multiplying it by the new correction map). For resting-state fMRI runs, independent component analysis and FMRIB's ICA-based Xnoisiefier (ICA+FIX) pipeline (*7*–*10*) were applied to resting-state fMRI scans to eliminate spatially specific temporally structured artifacts. The ICA+FIX pipeline consists of the following steps: 1) high-pass temporal filtering to eliminate linear trends in the data with a sigma of 1,000 s (run length = 864 s); 2) MELODIC ICA, creating component spatial maps and timeseries with auto-dimensionality selection of up to 250 components; (iii) the FIX-trained ICA component classifier categories these components into signal and noise; (iv) out-regression of the data and all ICA components of the 24 motion parameters (which were also temporal high-pass-filtered with a sigma of 1,000 s). All of the ICA component timeseries were utilized to calculate regression coefficients, and the noise component timeseries were weighted by the coefficients and subtracted from the data (a "non-aggressive" regression approach). The volumetric timeseries data was processed through the ICA+FIX algorithms before the grayordinates timeseries data underwent the high-pass filter and nuisance regression stages. After the original, uncleaned, native mesh data had been resampled into the standard grayordinates space in accordance with the areal feature-based MSM surface registration, the ICA+FIX cleanup was once again applied to the resting state fMRI-dense timeseries data. Early analyses included regression of the mean gray signal ("global signal"); however, this method was abandoned since it shifted some resting-state fMRI functional connectivity gradient locations, thereby, reducing cross-modal alignment. No further spatial smoothing or temporal low-pass filtering was carried out because these types of "lossy" preprocessing steps would decrease the accuracy of the parcellations and proved unnecessary for the purposes of the current study. Using GraphVar (*11*), we then developed participant-specific connection matrices from the timeseries signals of 360 brain regions. We employed the 360 areas that were defined by HCP-style parcellation (*12*, *13*) as nodes to build the brain network. Using Pearson's correlation coefficients, the edges of the brain network were defined as the functional connectivity of all pairs among the 360 areas.

HCP pipelines were also used to preprocess diffusion MRI data. Briefly, corrections for gradient, B0, eddy current distortions, and cross-modal registration were performed (*2*, *14*). The intensity was normalized by the mean of volumes with b = 0 s/mm$^2$ (b0 volumes), and two opposing phase-encoded images and FSL's Topup (*15*) were used to correct B0-inhomogeneity distortion. Prior to the recent re-computation of HCP, which included outlier detection (*16*), FSL's Eddy, version 5.0.9 was used to correct the eddy current-induced field inhomogeneities and head motion for each image volume. The data were then adjusted for gradient non-linearity. The b0 volume and the BBR cost function in FSL and FreeSurfer's BBRegister were used to register the diffusion data to the structural T1-weighted anterior commissure-posterior commissure space and the white matter surface, respectively. Based on the rotational information of the b0 to the T1-weighted transformation matrix, the diffusion gradient vectors were rotated.

To calculate the neurite orientation dispersion and density imaging (NODDI) coefficients, the AMICO toolbox was used. We modified the AMICO toolbox and changed its intrinsic free diffusivity parameter to $1.1 \times 10^{-3}$ mm$^2$/s for our analyses of gray matter structures because it is essential to optimize the NODDI model when analyzing gray matter structures because different types of brain tissue may vary considerably regarding their intrinsic free diffusivity. Subsequently, we calculated the NDI (the amount of stick-like or cylindrically symmetric diffusion produced when water molecules are constrained by neurite membranes) and ODI (a tortuosity measure, coupling the intra- and extra-neurite space, resulting in alignment or dispersion of axons and dendrites in the gray matter). Using NoddiSurfaceMapping (*17*), the NODDI parameters were mapped onto the cortical surface. Using FSL's bedpostX and probtrackX methods (*18*, *19*), probabilistic tractography was performed to obtain tractography matrix from the preprocessed diffusion MRI data. To obtain a tractography matrix of the number of streamlines originating from each regions-of-interest (ROI) and reaching the rest of the cerebral cortex, we seeded 1,000 streamlines from each of the ROIs. By dividing each row by the waytotal file, the unnormalized values in these matrices were normalized.

We calculated the nodal graph measures using resting-state functional connectivity and tract based structural connectivity matrix. Thresholding of the connectivity matrix was done. To find the optimal threshold value, we performed a graph theory-based analysis (*20*) seeking a threshold value that maximizes the global cost efficiency. We used segregation (clustering coefficient and local efficiency), integration (nodal path length), and centrality (degree and betweenness centrality) measures, that most commonly used concept for describing the topological property of a node in a network. These nodal graph measures were calculated using GraphVar (*21*).

Statistical analysis

All statistical analyses were conducted using R Studio (version 1.1.463). All variables were regressed from age and sex before analyses. Brain imaging variables were additionally regressed from handedness, intracranial volume, and transmitter reference amplitude. Multiple sparse canonical correlation analysis (MSCCA) was performed using the MultiCCA.permute and MultiCC functions in the PMA package to test the association of prosocial behavior and imaging data. Optimal weights and penalties were identified using the MultiCCA.permute function with 1,000 permutations. Before performing MSCCA, principal component analysis (PCA) was performed for dimensionality reduction in all sets of variables (i.e., economic games, cortical and subcortical structure, resting-state functional connectivity, and tractography-based structural connectivity) using the prcomp function. A dimensionality reduction step was performed to prevent an overdetermined, rank-deficient solution and eliminate the possibility of overfitting. The data obtained from economic games were reduced to 30 PCAs (variance explained = 89%) and all sets of MRI variables were reduced to 60 PCAs (variance explained = 64 to 72%). The permutation test confirmed whether the sum of canonical correlations was statistically significant relative to the null distribution. To produce the null distribution, the PCAs were randomly shuffled 1,000 times before performing MSCCA. The $p$-value was calculated as the proportion of cases that showed a higher canonical correlation than the observed correlation, which was divided by 1,000. The results of permutation test are shown in **Fig. S1**. To confirm the robustness of MSCCA when varying the number of principal components, we repeated the whole statistical testing for a range of PCs (i.e., 70 and 80) for MRI variables as supplementary analyses. The results of these analyses are shown in **Figs. S4** and **S5**.

Supplementary analysis for robustness of the MSCCA

The results of canonical cross-loadings for prosocial behavior and brain imaging data with variation in the number of principal components are presented in **Fig. S4**. The canonical cross-loading for economic games and brain imaging data were robust with variation in the number of principal components. The correlation among the canonical cross-loadings for brain imaging data in each index when varying the number of principal components is shown in **Fig. S5**. There was almost no change in the results—the correlations are [0.76 to 0.99].
