## Supplementary material for "Multimodal imaging for identifying brain markers of human prosocial behavior": Figs. S1 to S5

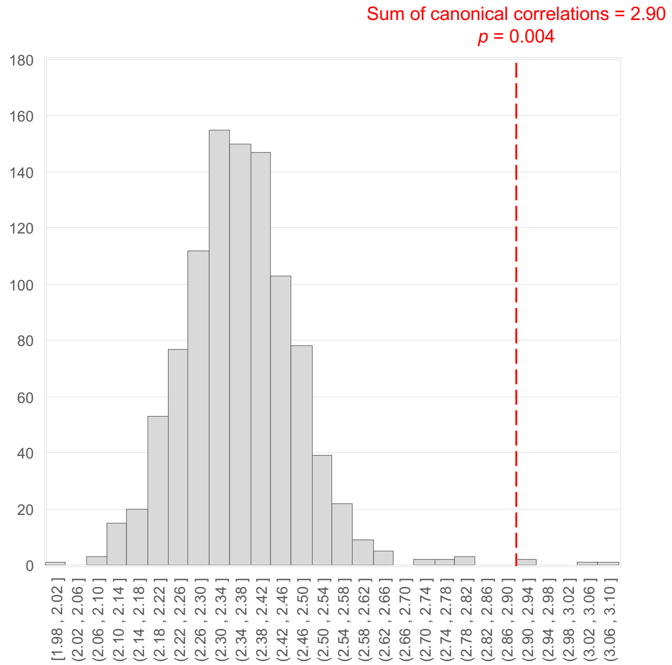


**Fig. S1.** The result of the permutation test for the multiple sparse canonical correlation analysis. The null distribution of canonical correlations with randomly shuffled data (histogram) and the true canonical correlation (red dashed line).

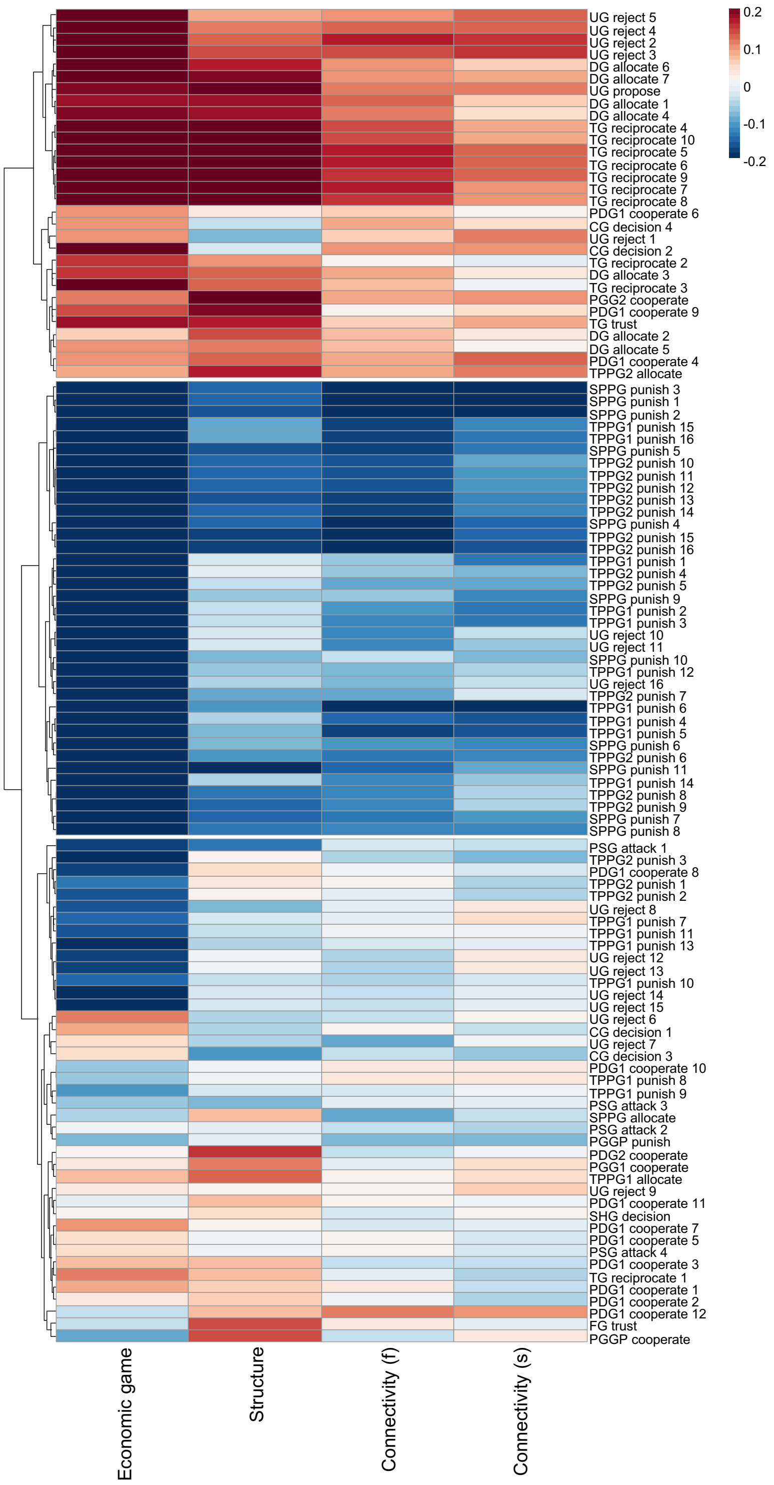


**Fig. S2.** Complete list of canonical cross-loadings for prosocial behaviors. Descriptions of the variables measured in each economic game are presented in **Table S2**.

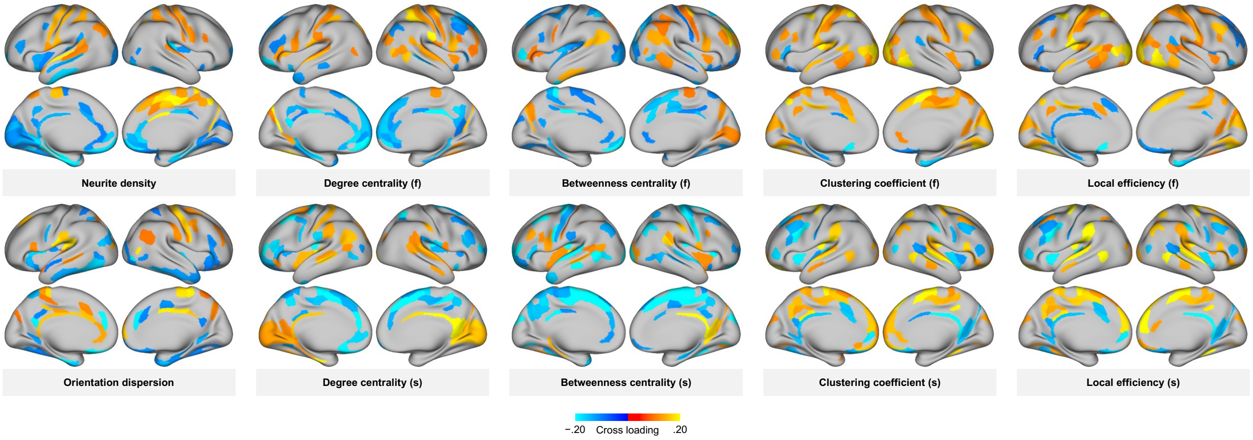


**Fig. S3.** A brain regional pattern of the strong covariation of brain imaging data with the economic games. This figure maps brain regions that show only the top 30% canonical correlation coefficients to confirm the brain regions strongly associated with prosociality. (f) denotes measures of resting-state functional connectivity; (s) denotes measures of tractography-based structural connectivity.

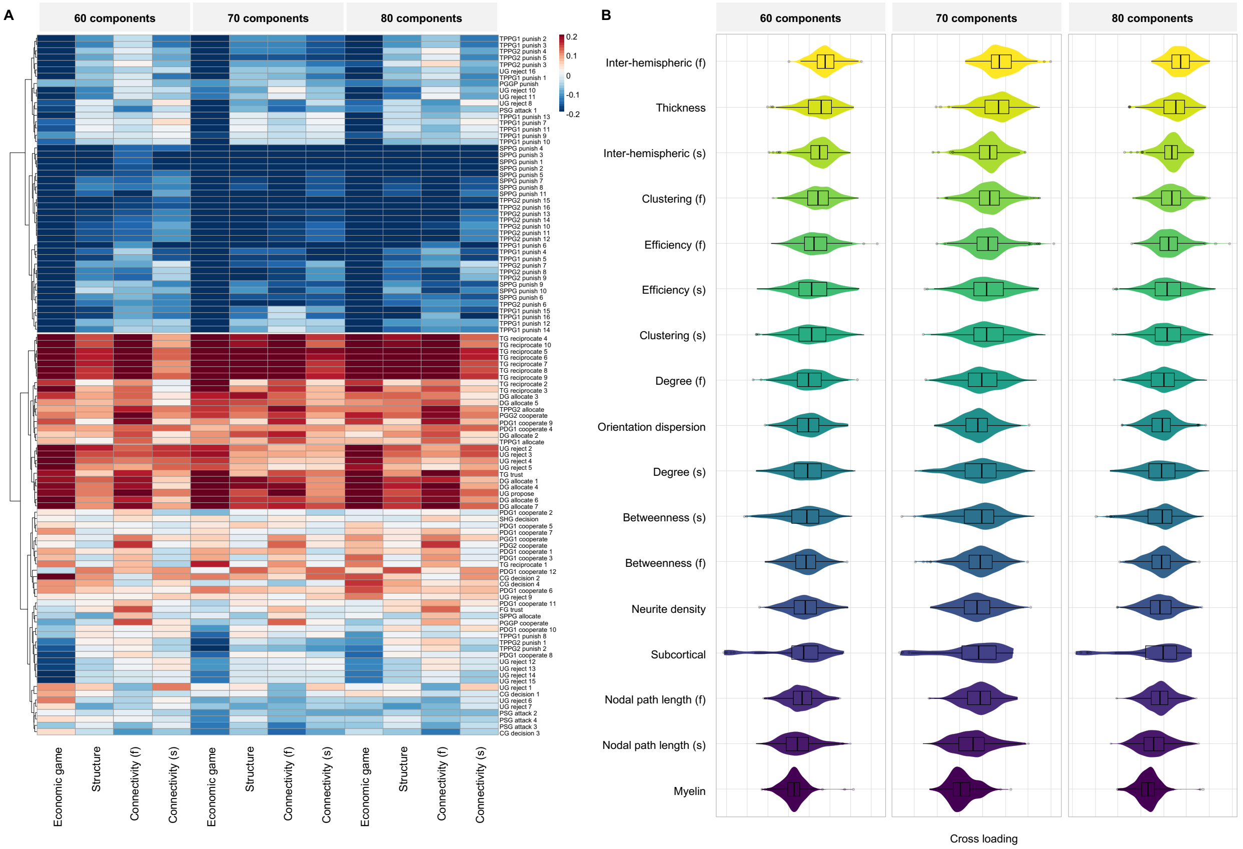


**Fig. S4.** Results of canonical cross-loadings for prosocial behavior and brain imaging data with variation in the number of principal components. (**A**) Canonical cross-loadings for prosocial behavior, (**B**) Mean canonical cross-loadings for brain imaging data. Descriptions of the variables measured in each economic game are presented in **Table S2**.

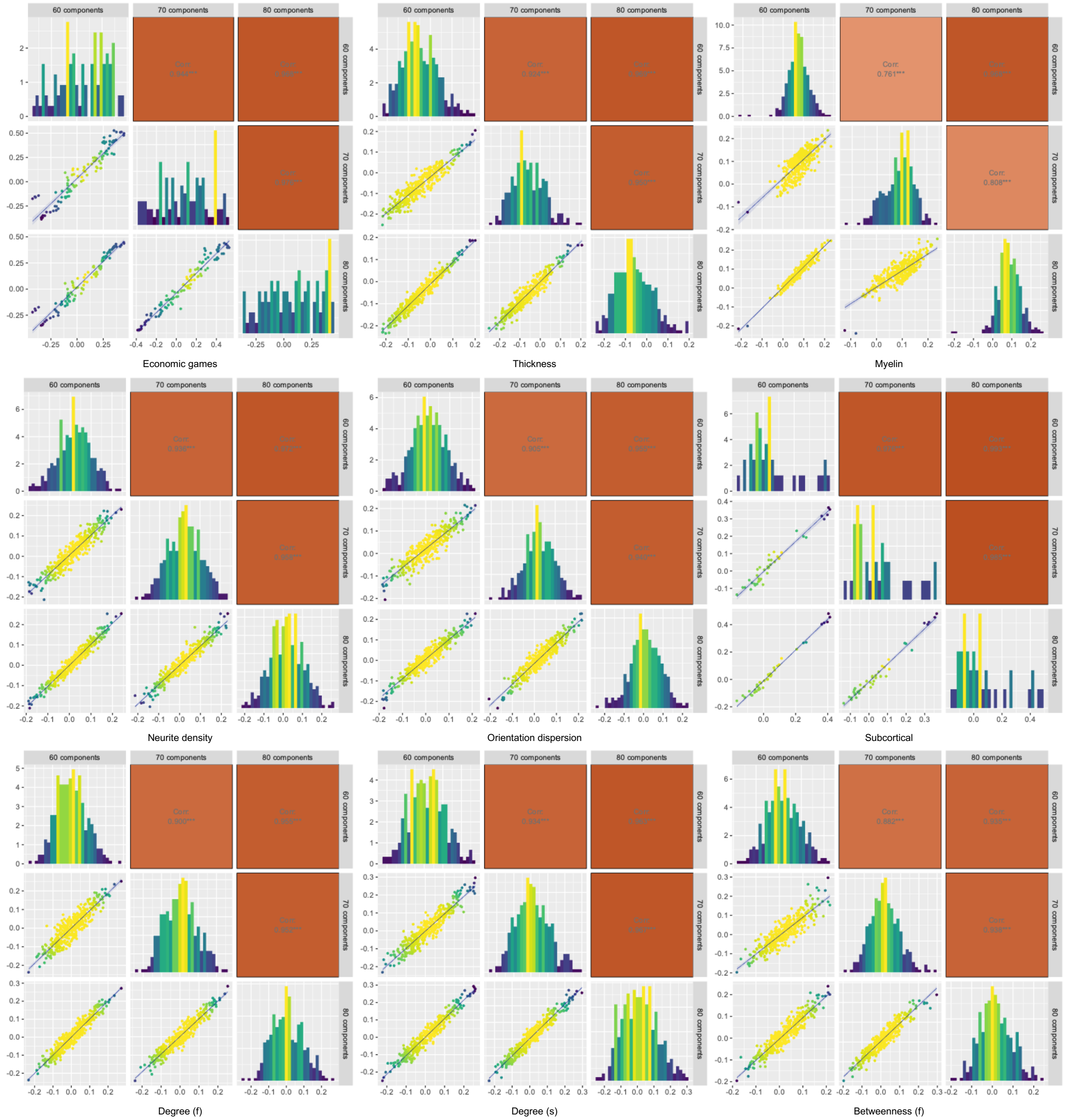
60 components
70 components
80 components
Economic games
Thickness
Myelin
Corr:
0.761***
0.808***
Neurite density
Orientation dispersion
Subcortical
Degree (f)
Degree (s)
Betweenness (f)
0.882***

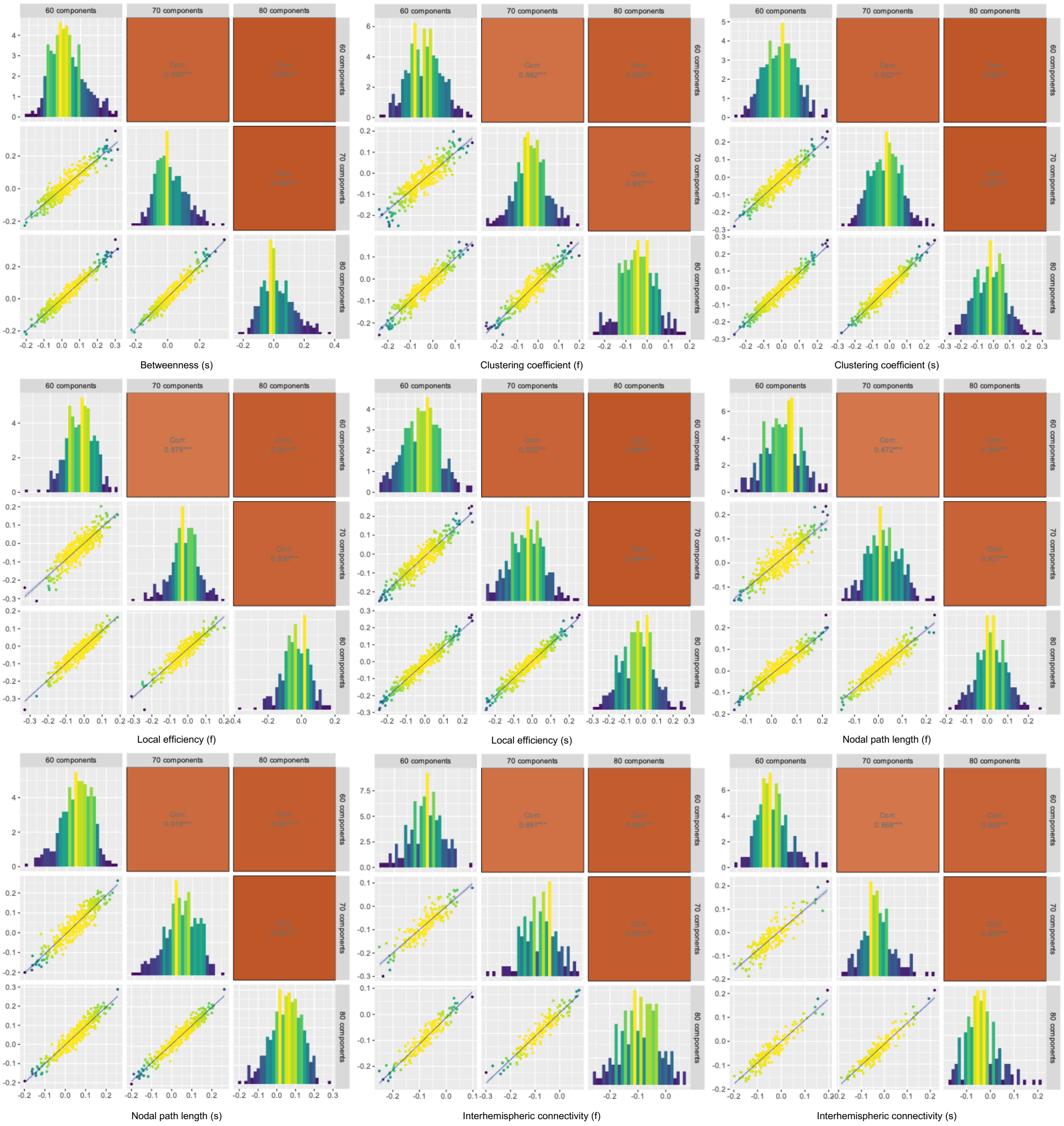


**Fig. S5.** Correlation among canonical cross-loadings for brain imaging data in each index with variation in the number of principal components
