## Supplementary material for "Multimodal imaging for identifying brain markers of human prosocial behavior": Tables S1 to S4

**Table S1.**
Complete list of canonical loadings.

Due to the large amount of data, **Table S1** is stored in an external repository.
https://osf.io/r2ewv/?view_only=f6bb9247afa74713a23337a24c280435

**Table S2.**

Description of variables used in the analysis.

| Variable | Description |
|---|---|
| Prisoner's Dilemma Game (PDG) I | |
| PDG1 cooperate 1 | The decision under condition simultaneous PDG (0 = not provided, 1 = provided). The endowment is JPY 300. |
| PDG1 cooperate 2 | The decision under condition simultaneous PDG (0 = not provided, 1 = provided). The endowment is JPY 800. |
| PDG1 cooperate 3 | The decision under condition simultaneous PDG (0 = not provided, 1 = provided). The endowment is JPY 1500. |
| PDG1 cooperate 4 | The decision under condition sequential PDG first (0 = not provided, 1 = provided). The endowment is JPY 300. |
| PDG1 cooperate 5 | The decision under condition sequential PDG first (0 = not provided, 1 = provided). The endowment is JPY 800. |
| PDG1 cooperate 6 | The decision under condition sequential PDG first (0 = not provided, 1 = provided). The endowment is JPY 1500. |
| PDG1 cooperate 7 | Under condition sequential PDG second, the decision when the opponent provide (0 = not provided, 1 = provided). The endowment is JPY 300. |
| PDG1 cooperate 8 | Under condition sequential PDG second, the decision when the opponent did not provide (0 = not provided, 1 = provided). The endowment is JPY 300. |
| PDG1 cooperate 9 | Under condition sequential PDG second, the decision when the opponent provide (0 = not provided, 1 = provided). The endowment is JPY 800. |
| PDG1 cooperate 10 | Under condition sequential PDG second, the decision when the opponent did not provide (0 = not provided, 1 = provided). The endowment is JPY 800. |
| PDG1 cooperate 11 | Under condition sequential PDG second, the decision when the opponent provide (0 = not provided, 1 = provided). The endowment is JPY 1500. |
| PDG1 cooperate 12 | Under condition sequential PDG second, the decision when the opponent did not provide (0 = not provided, 1 = provided). The endowment is JPY 1500. |
| Dictator Game (DG) | |
| DG allocate 1 | The rate of distribution at one-shot DG. The endowment is JPY 1000. |
| DG allocate 2 | The amount of distribution rate at repeated one-shot DG. The endowment is JPY 300. |
| DG allocate 3 | The amount of distribution rate at repeated one-shot DG. The endowment is JPY 400. |
| DG allocate 4 | The amount of distribution rate at repeated one-shot DG. The endowment is JPY 600. |
| DG allocate 5 | The amount of distribution rate at repeated one-shot DG. The endowment is JPY 700. |

| | |
|---|---|
| DG allocate 6 | The amount of distribution rate at repeated one-shot DG. The endowment is JPY 1200. |
| DG allocate 7 | The amount of distribution rate at repeated one-shot DG. The endowment is JPY 1300. |
| Faith Game (FG) | |
| FG trust | The rate of the amount left to the distributor |
| Public Goods Game (PGG) I | |
| PGG1 cooperate | The rate of provision to the group |
| Prisoner's Dilemma Game (PDG) II | |
| PDG2 cooperate | The rate of JPY 1000 provided to the opponent |
| Second-party punishment game (SPPG) | |
| SPPG allocate | The rate of provision in PDG with punishment |
| SPPG punish 1 | The amount of money spent in punitive PDG when the opponent provided JPY 0. |
| SPPG punish 2 | The amount of money spent in punitive PDG when the opponent provided JPY 100. |
| SPPG punish 3 | The amount of money spent in punitive PDG when the opponent provided JPY 200. |
| SPPG punish 4 | The amount of money spent in punitive PDG when the opponent provided JPY 300. |
| SPPG punish 5 | The amount of money spent in punitive PDG when the opponent provided JPY 400. |
| SPPG punish 6 | The amount of money spent in punitive PDG when the opponent provided JPY 500. |
| SPPG punish 7 | The amount of money spent in punitive PDG when the opponent provided JPY 600. |
| SPPG punish 8 | The amount of money spent in punitive PDG when the opponent provided JPY 700. |
| SPPG punish 9 | The amount of money spent in punitive PDG when the opponent provided JPY 800. |
| SPPG punish 10 | The amount of money spent in punitive PDG when the opponent provided JPY 900. |
| SPPG punish 11 | The amount of money spent in punitive PDG when the opponent provided JPY 1000. |
| Trust Game (TG) | |
| TG trust | The transfer rate by trustor |
| TG reciprocate 1 | The amount returned by trustee if trustor transfers JPY 100 to trustee |
| TG reciprocate 2 | The amount returned by trustee if trustor transfers JPY 200 to trustee |
| TG reciprocate 3 | The amount returned by trustee if trustor transfers JPY 300 to trustee |
| TG reciprocate 4 | The amount returned by trustee if trustor transfers JPY 400 to trustee |

| | |
| --- | --- |
| TG reciprocate 5 | The amount returned by trustee if trustor transfers JPY 500 to trustee |
| TG reciprocate 6 | The amount returned by trustee if trustor transfers JPY 600 to trustee |
| TG reciprocate 7 | The amount returned by trustee if trustor transfers JPY 700 to trustee |
| TG reciprocate 8 | The amount returned by trustee if trustor transfers JPY 800 to trustee |
| TG reciprocate 9 | The amount returned by trustee if trustor transfers JPY 900 to trustee |
| TG reciprocate 10 | The amount returned by trustee if trustor transfers JPY 1000 to trustee |
| Ultimatum Game (UG) | |
| UG propose | The rate offered by proposer |
| UG reject 1 | The decisions made by responders when offering JPY 0 (0 = accept, 1 = reject) |
| UG reject 2 | The decisions made by responders when offering JPY 100 (0 = accept, 1 = reject) |
| UG reject 3 | The decisions made by responders when offering JPY 200 (0 = accept, 1 = reject) |
| UG reject 4 | The decisions made by responders when offering JPY 300 yen (0 = accept, 1 = reject) |
| UG reject 5 | The decisions made by responders when offering JPY 400 yen (0 = accept, 1 = reject) |
| UG reject 6 | The decisions made by responders when offering JPY 500 yen (0 = accept, 1 = reject) |
| UG reject 7 | The decisions made by responders when offering JPY 600 yen (0 = accept, 1 = reject) |
| UG reject 8 | The decisions made by responders when offering JPY 700 yen (0 = accept, 1 = reject) |
| UG reject 9 | The decisions made by responders when offering JPY 800 yen (0 = accept, 1 = reject) |
| UG reject 10 | The decisions made by responders when offering JPY 900 yen (0 = accept, 1 = reject) |
| UG reject 11 | The decisions made by responders when offering JPY 1000 yen (0 = accept, 1 = reject) |
| UG reject 12 | The decisions made by responders when offering JPY 1100 yen (0 = accept, 1 = reject) |
| UG reject 13 | The decisions made by responders when offering JPY 1200 yen (0 = accept, 1 = reject) |
| UG reject 14 | The decisions made by responders when offering JPY 1300 yen (0 = accept, 1 = reject) |
| UG reject 15 | The decisions made by responders when offering JPY 1400 yen (0 = accept, 1 = reject) |
| UG reject 16 | The decisions made by responders when offering JPY 1500 yen (0 = accept, 1 = reject) |
| Third-party Punishment Game (TPPG) I | |

| | |
|---|---|
| TPPG1 punish 1 | The amount of penalty by a third-party if the distributor distributes JPY 0 to the recipient |
| TPPG1 punish 2 | The amount of penalty by a third-party if the distributor distributes JPY 100 to the recipient |
| TPPG1 punish 3 | The amount of penalty by a third-party if the distributor distributes JPY 200 to the recipient |
| TPPG1 punish 4 | The amount of penalty by a third-party if the distributor distributes JPY 300 to the recipient |
| TPPG1 punish 5 | The amount of penalty by a third-party if the distributor distributes JPY 400 to the recipient |
| TPPG1 punish 6 | The amount of penalty by a third-party if the distributor distributes JPY 500 to the recipient |
| TPPG1 punish 7 | The amount of penalty by a third-party if the distributor distributes JPY 600 to the recipient |
| TPPG1 punish 8 | The amount of penalty by a third-party if the distributor distributes JPY 700 to the recipient |
| TPPG1 punish 9 | The amount of penalty by a third-party if the distributor distributes JPY 800 to the recipient |
| TPPG1 punish 10 | The amount of penalty by a third-party if the distributor distributes JPY 900 to the recipient |
| TPPG1 punish 11 | The amount of penalty by a third-party if the distributor distributes JPY 1000 to the recipient |
| TPPG1 punish 12 | The amount of penalty by a third-party if the distributor distributes JPY 1100 to the recipient |
| TPPG1 punish 13 | The amount of penalty by a third-party if the distributor distributes JPY 1200 to the recipient |
| TPPG1 punish 14 | The amount of penalty by a third-party if the distributor distributes JPY 1300 to the recipient |
| TPPG1 punish 15 | The amount of penalty by a third-party if the distributor distributes JPY 1400 to the recipient |
| TPPG1 punish 16 | The amount of penalty by a third-party if the distributor distributes JPY 1500 to the recipient |
| TPPG1 allocate | The distribution rate by distributor |
| Third-party Punishment Game II | |
| TPPG2 punish 1 | The amount of penalty by a third-party if the distributor distributes JPY 0 to the recipient |
| TPPG2 punish 2 | The amount of penalty by a third-party if the distributor distributes JPY 100 to the recipient |
| TPPG2 punish 3 | The amount of penalty by a third-party if the distributor distributes JPY 200 to the recipient |

| | |
|---|---|
| TPPG2 punish 4 | The amount of penalty by a third-party if the distributor distributes JPY 300 to the recipient |
| TPPG2 punish 5 | The amount of penalty by a third-party if the distributor distributes JPY 400 to the recipient |
| TPPG2 punish 6 | The amount of penalty by a third-party if the distributor distributes JPY 500 to the recipient |
| TPPG2 punish 7 | The amount of penalty by a third-party if the distributor distributes JPY 600 to the recipient |
| TPPG2 punish 8 | The amount of penalty by a third-party if the distributor distributes JPY 700 to the recipient |
| TPPG2 punish 9 | The amount of penalty by a third-party if the distributor distributes JPY 800 to the recipient |
| TPPG2 punish 10 | The amount of penalty by a third-party if the distributor distributes JPY 900 to the recipient |
| TPPG2 punish 11 | The amount of penalty by a third-party if the distributor distributes JPY 1000 to the recipient |
| TPPG2 punish 12 | The amount of penalty by a third-party if the distributor distributes JPY 1100 to the recipient |
| TPPG2 punish 13 | The amount of penalty by a third-party if the distributor distributes JPY 1200 to the recipient |
| TPPG2 punish 14 | The amount of penalty by a third-party if the distributor distributes JPY 1300 to the recipient |
| TPPG2 punish 15 | The amount of penalty by a third-party if the distributor distributes JPY 1400 to the recipient |
| TPPG2 punish 16 | The amount of penalty by a third-party if the distributor distributes JPY 1500 to the recipient |
| TPPG2 allocate | The distribution rate by distributor |
| Preemptive strike game (PSG) | |
| PSG attack 1 | Whether the button was pressed in condition 55 (o = not pressed, 1 = pressed) |
| PSG attack 2 | Whether the button was pressed in condition 1010 (o = not pressed, 1 = pressed) |
| PSG attack 3 | Whether the button was pressed in condition 510 (o = not pressed, 1 = pressed) |
| PSG attack 4 | Whether the button was pressed in condition 105 (o = not pressed, 1 = pressed) |
| Chicken Game (CG) | |
| CG decision 1 | The first decision with simultaneous conditions (0 = turn back, 1 = proceed) |
| CG decision 2 | The second decision with simultaneous conditions (0 = turn back, 1 = proceed) |

| | |
|---|---|
| CG decision 3 | The third decision with simultaneous conditions (0 = turn back, 1 = proceed) |
| CG decision 4 | The forth decision with participant first condition (0 = turn back, 1 = proceed) |
| Public Goods Game (PGG) II | |
| PGG2 cooperate | Rate of contribution in the Public Goods Game without punishment |
| Public Goods Game with punishment (PGGP) | |
| PGGP cooperate | Rate of contribution in the Public Goods Game with punishment |
| PGGP punish | Amounts used for the punishment system |
| Stag Hunt Game (SHG) | |
| SHG decision | Decision to invest or not (0 = keep, 1 = invest) |

**Table S3.**

Schedule for collecting the data used in this study

| |
|---|
| **Wave 1:** n = 564, Day = May 2012 – July 2012 |
| Demographic data |
| **Wave 2**: n = 483, Day = October 2012 – February 2013 |
| Prisoner's dilemma game I (12 variables) |
| **Wave 3**: n = 489, Day = April 2013 – June 2013 |
| Dictator game (7 variables) |
| Faith game (1 variables) |
| **Wave 4**: n = 474, Day = September 2013 – October 2013 |
| Public goods game I (1 variable) |
| Prisoner's dilemma game II (1 variable) |
| Second-party punishment game (12 variables) |
| **Wave 5**: n = 471, Day = December 2013 – February 2014 |
| Trust game (11 variables) |
| Ultimatum game (17 variables) |
| **Wave 6**: n = 470, Day = May 2014 – July 2014 |
| Third-party punishment game I (17 variables) |
| Third-party punishment game II (17 variables) |
| Preemptive strike game (4 variables) |
| **Wave 7**: n = 451, Day = October 2014 – January 2015 |
| Chicken game (4 variables) |
| **Wave 8**: n = 424, Day = September 2015 – December 2015 |
| Public Goods Game II (1 variable) |
| Public Goods Game with punishment (2 variables) |
| Stag Hunt Game (1 variable) |
| **Wave 9**: n = 290, Day = November 2016 – March 2018 |
| MRI data (T1w, T2w, resting-state fMRI, diffusion MRI) |
| **Wave 10**: n = 307, Day = July 2018 – November 2018 |

fMRI, functional MRI; MRI, magnetic resonance imaging

**Table S4.**

List of papers published in this research project

| No | Bibliographic information of the article |
|---|---|
| 1 | H. Tanaka, Q. Shou, T. Kiyonari, T. Matsuda, M. Sakagami, H. Takagishi, Right dorsolateral prefrontal cortex regulates default prosociality preference. *Cerebral Cortex* **33**, 5420-5425 (2023). |
| 2 | A.S.R. Fermin, T. Kiyonari, Y. Matsumoto, H. Takagishi, Y. Li, R. Kanai, M. Sakagami, R. Akaishi, N. Ichikawa, M. Takamura, S. Yokoyama, M.G. Machizawa, H.L. Chan, A. Matani, S. Yamawaki, G. Okada, Y. Okamoto, T. Yamagishi, The neuroanatomy of social trust predicts depression vulnerability. *Scientific Reports* **12**, 16724 (2022). |
| 3 | Q. Shou, J. Yamada, K. Nishina, M. Matsunaga, T. Matsuda, H. Takagishi, Association between salivary oxytocin levels and the amygdala and hippocampal volumes. *Brain Structure and Function* **227**, 2503–2511 (2022). |
| 4 | Q. Shou, J. Yamada, K. Nishina, M. Matsunaga, T. Kiyonari, H. Takagishi, Is oxytocin a trust hormone? Salivary oxytocin is associated with caution but not with general trust. *PLOS ONE* **17**, e0267988 (2022). |
| 5 | K. Nishina, Q. Shou, H. Takahashi, M. Sakagami, M. Inoue-Murayama, H. Takagishi, Association between polymorphism (5-HTTLPR) of the serotonin transporter gene and behavioral response to unfair distribution. *Frontiers in Behavioral Neuroscience* **16**, 762092 (2022). |
| 6 | J. Yamada, Y. Nakawake, Q. Shou, K. Nishina, M. Matsunaga, H. Takagishi, Salivary oxytocin is negatively associated with religious faith in Japanese non-Abrahamic people. *Frontiers in Psychology* **12**, 705781 (2012). |
| 7 | T. Ishihara, A. Miyazaki, H. Tanaka, T. Fujii, M. Takahashi, K. Nishina, K. Kanari, H. Takagishi, T. Matsuda, Childhood exercise predicts response inhibition in later life via changes in brain connectivity and structure. *NeuroImage* **237**, 118196 (2021). |
| 8 | K. Nishina, H. Takagishi, H. Takahashi, M. Sakagami, M. Inoue-Murayama, Association of polymorphism of arginine-vasopressin receptor 1A (AVPR1a) gene with trust and reciprocity. *Frontiers in Human Neuroscience* **13**, 230 (2019). |
| 9 | K. Nishina, H. Takagishi, A.S.R. Fermin, M. Inoue-Murayama, H. Takahashi, M. Sakagami, T. Yamagishi, Association of the oxytocin receptor gene with attitudinal trust: role of amygdala volume. *Social Cognitive and Affective Neuroscience* **13**, 1091–1097 (2018). |
| 10 | T. Yamagishi, Y. Li, A.S.R. Fermin, R. Kanai, H. Takagishi, Y. Matsumoto, T. Kiyonari, M. Sakagami, Behavioural differences and neural substrates of altruistic and spiteful punishment. *Scientific Reports* **7**, 14654 (2017). |
| 11 | T. Yamagishi, Y. Matsumoto, T. Kiyonari, H. Takagishi, Y. Li, R. Kanai, M. Sakagami, Response time in economic games reflects different types of decision conflict for prosocial and proself individuals. *Proceedings of the National Academy of Sciences of the United States of America* **114**, 6394–6399 (2017). |
| 12 | T. Yamagishi, H. Takagishi, A.S.R. Fermin, R. Kanai, Y. Li, Y. Matsumoto, Cortical thickness of the dorsolateral prefrontal cortex predicts strategic choices in economic games. *Proceedings of the National Academy of Sciences of the United States of America* **113**, 5582–5587 (2016). |
| 13 | Y. Matsumoto, T. Yamagishi, Y. Li, T. Kiyonari, Prosocial behavior increases with age across five economic games. *PLOS ONE* **11**, e0158671 (2016). |

14 T. Yamagishi, Y. Li, Y. Matsumoto, T. Kiyonari, Moral bargain hunters purchase moral righteousness when it is cheap: within-individual effect of stake size in economic games. *Scientific Reports* **6**, 27824 (2016).
15 K. Nishina, H. Takagishi, M. Inoue-Murayama, H. Takahashi, T. Yamagishi, Polymorphism of the Oxytocin Receptor Gene Modulates Behavioral and Attitudinal Trust among Men but Not Women. *PLOS ONE* **10**, e0137089 (2015).
16 T. Yamagishi, Y. Li, H. Takagishi, Y. Matsumoto, T. Kiyonari, In search of homo economicus. *Psychological science* **25**, 1699–1711 (2014).
